# Microbiological Survey of the San Pedro Basin Subseafloor

**DOI:** 10.64898/2026.01.28.702002

**Authors:** Dyanna Jimenez, Fernando Paz-Melchor, Xiuhnel Zien, Tiffany Smith, Gisselle Camas, Nicolina Trejo, Shiva Sadeghpour, M. Oscar Almacen, Seshan G. Goluhewa, Arkaidy I. Garber, Lilian Real-Nuñez, Gustavo A. Ramírez

## Abstract

Marine subseafloor sediments host extensive microbial communities that drive global biogeochemical processes. Despite its proximity to one of the most densely populated and economically important coastal regions in the world, the sub benthic ecology of the San Pedro Channel, the waterway separating Los Angeles County from Santa Catalina Island, remains largely uncharacterized. To establish the foundational knowledge required for future impact assessment studies, we initiated a sediment coring survey that generated a water and sediment depth standardized transect across the sloping flanks of the San Pedro Basin. Here, we describe the early development of an integrated subseafloor microbiological catalogue for this region of intense maritime activity incorporating molecular ecology, microbial isolation and cultivation, and physiological assays. This initial effort is designed not as a direct comparison between basin flanks, but as a baseline assessment that captures ecological and microbiological variation across paired sides matched in water column depth and sediment depth beneath seafloor. Accordingly, we provide a cross channel coordinated dataset of subseafloor microbial communities and cultivated representatives from San Pedro Basin sediment, offering a critical starting point for understanding natural variability and for detecting potential signatures of environmental or anthropogenic disturbance. Ultimately, this baseline will support long□term monitoring efforts and supply a curated collection of isolates for future experimental microbiology and comparative genomics research.

## Introduction

The subseafloor is one of Earth’s largest microbial habitats, with coastal sediments hosting the highest cell densities and collectively containing more microbial biomass than ocean water or terrestrial soils (Kallmeyer et al. 2012; Bar-On, Phillips, and Milo 2018). Coastal sediments are enriched in organic carbon and can contain over a billion prokaryotic cells per cubic centimeter (Kallmeyer et al. 2012; Hoshino and Inagaki 2019). These taxonomically diverse and metabolically complex microbial communities drive biogeochemical cycles on timescales ranging from decades to hundreds of millions of years (D’Hondt et al. 2004; D’Hondt et al. 2019). Due to slow burial (Jorgensen and Marshall 2016), diffusion-limited oxidant and product transport (D’Hondt et al. 2019), and persistent electron acceptor scarcity (Bradley et al. 2020) subseafloor sediments are energy-limited environments. Despite this, microbes persists in the global subseafloor over geological timescales (Morono et al. 2020) and significantly influence major nutrient cycling (C, N, S), trace metal bio-availability (Fe, Mn, Ni), and the generation of seafloor materials of economic interest (*e.g.* hydrocarbon reserves).

The San Pedro Channel is a maritime corridor bounded by the steep slopes of the mainland San Pedro shelf and Santa Catalina Island and an area of global economic importance (Figure 1). Beneath it, the San Pedro Basin, bound by the Palos Verdes margin and Santa Catalina ridge, is a tectonically active marine basin characterized by subsidence and steep margins (Normark et al. 2009). The eastern basin margin traps terrigenous material from the Los Angeles coastal plain as riverine input and mass wasting events (Normark et al. 2009; Warrick and Farnsworth 2009). The western margin, descending form Santa Catalina Island, exhibits lower sedimentation rates, greater stratigraphic continuity, and more stable depositional conditions dominated by hemipelagic processes (Fisher et al. 2004). Uneven accumulation, due to internal tide generation (Garrett and Kunze 2007), results in heterogenous stratigraphy generating mixed-age deposits and variable sedimentation rates across short spatial scales in the region (Fisher et al. 2004; Normark et al. 2009).

**Figure 1.**
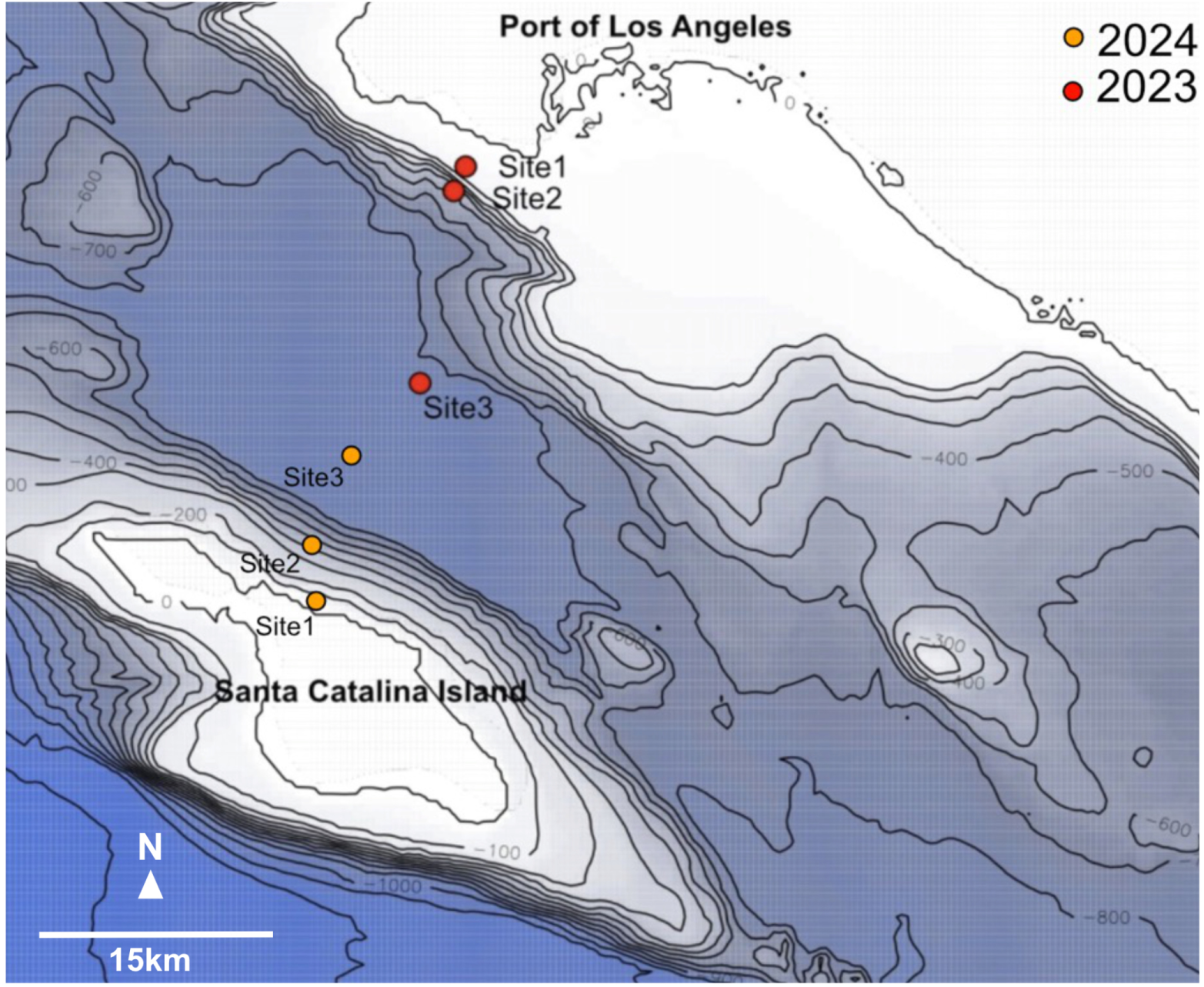
Bathymetric map of the San Pedro Basin located between Santa Catalina Island and Los Angeles in the California Bight region of the Borderlands. At both channel slopes, sites represent the following water depths: Site 1: 65 m, Site 2: 401 m, Site 3: 893 m. Site color depicts collection year with red and orange for the 2023 and 2024 R/V Yellowfin coring campaigns, respectively.

Sediment cores function as natural time capsules capturing vertical ecological information that preserve markers of microbial community structure, metabolic activity, and environmental change (Kirkpatrick, Walsh, and D’Hondt 2019; Morono et al. 2009; Orsi, Magritsch, Vargas, Coskun, Vuillemin, Höhna, Wörheide, D’Hondt, et al. 2021). Where sedimentation rate models exists, cores allow reconstruction of historical environmental dynamics, enabling inferences about both natural variability and anthropogenic disturbance in aquatic systems (Valette-Silver 1993). For example, in limnetic and freshwater systems, sediment microbiome analyses are routinely used to trace environmental change, eutrophication, pollution, and industrial impacts (Miller 2014; Mills 2017; Simon 2023).

In the San Pedro Channel region, through stormwater runoff (Ahn and Grant 2007), shipping activity (Xiao et al. 2022), legacy industrial impacts (Eganhouse and Pontolillo 2000), and the discharge of an estimated 15–30 million gallons of treated wastewater per day (California Regional Water Quality Control Board 2016), the Los Angeles metropolitan area disproportionally influences the eastern basin ecosystem. Together, these inputs likely shape the ecology, resilience, and biogeochemical roles of subseafloor microbial communities and their activities. Nearly two decades ago, an automated rRNA intergenic spacer analysis (ARISA) survey of surficial marine sediment (top 1 cm only) across opposite sides of the San Pedro Channel showed that shallow water sediment communities are similar across the channel while deeper water surficial sediments exhibit weak taxon coupling and strong environmental structuring (Hewson, Jacobson Meyers, and Fuhrman 2007). The study concludes that, despite the relative stability of abundance and diversity metrics in the sediment interface across the region, taxon composition can change dramatically suggesting the need for higher taxon resolution and boarder spatial coverage, to better understand dispersal, selection, and function in the regional subseafloor microbial ecosystem.

We report the collection of sediment cores along water depth-matched transects across the Santa Catalina and Los Angeles slopes of the San Pedro Basin. We used these samples, spanning full-channel water depth, to generate ecological, physiological, and comparative genomics analyses of subseafloor sediment microbiomes. These samples form the basis of an emerging molecular and microbiological dataset that may eventually evaluate ecological impacts associated with environmental pressures posed by a major coastal metropolis on the microbial ecology of the hidden, yet nearby, subseafloor.

## Materials and Methods

### Microbiological sampling

San Pedro channel sediment samples were collected using a gravity corer on the Southern California Marine Institute’s (SCMI) Research Vessel (RV) Yellowfin, as detailed elsewhere (Ramírez et al. 2018). Briefly, once on deck, duplicate 10-cc syringe mini-core samples were aseptically collected from the middle of the core liner using sterile technique. A 1-cc sample from a mini core was immediately used to generate a 1:10 slurry using sterilized artificial seawater. A 0.1 ml volume of fresh (within 10 mins, never frozen to avoid inadvertent cellular lysis) slurry was used to inoculate Zobel Marine Agar plates, as detailed below. The remaining sediment was immediately frozen at sea (-20°C on the R/V Yellowfin laboratory freezer) and same-day transferred to -80°C back on shore, for shore-based analysis.

### Environmental DNA Extraction and Sequencing

DNA extractions from frozen minicores and, after weeks of incubations, culture isolates were performed using FastDNA SPIN Kit for Soil (MP Biomedicals) as detailed elsewhere (Ramírez, Graham, and D’Hondt 2018). Briefly, this commercial kit protocol involves automatic homogenization in detergent-infused buffers for cell lysing, protein precipitation, silica-slurry binding of nucleic acids, and a final elution step using ultrapure water. Following DNA extraction, extract quantification was performed on a Nanodrop spectrophotometer. High-quality DNA extracts were outsourced for 16 rRNA gene sequencing, using the revised Earth Microbiome primer set (Parada, Needham, and Fuhrman 2016), on an Illumina platform at a depth of 30K reads per sample at Molecular Research DNA labs, Lubbock, Texas.

### 16S rRNA Gene Sequence Processing

Amplicon gene sequences were analyzed on R using the python SnakeMake workflow manager. Briefly, the DADA2 (Callahan et al. 2016) software was used for high-quality read assessments, paired-end read merging, and chimeric sequence screening to generate an error model used for amplicon sequence variants (ASVs) generation. A tabulated ASVs summary was subsequently used for all downstream ecological analyses as encouraged elsewhere (Callahan, McMurdie, and Holmes 2017). Sequence taxonomy was assigned using the SILVA 138 database (Quast et al. 2013). DECONTAM was used to remove potential kit contaminants from the initial ASVs table using a p=0.05 abundance-based statistical threshold (Davis et al. 2018). Decontaminated tabular products, specifically the ASV counts and taxonomy tables, were merged with all available metadata to generate an S4 object used for all computational analysis implemented on the Phyloseq R package (McMurdie and Holmes 2013), as detailed below. All sequence data generated has been deposited in NCBI SRA under accession numbers: xxxx-xxxx.

### Ecological Analyses and Modeling

All statistical analyses including diversity estimates [*e.g.*: Shannon diversity metric (Shannon and Weaver 1949)], multivariate modeling and visualization were generated in Phyloseq using the R-package Vegan (Oksanen et al. 2024). Specifically, following best practices for multiscale microbial ecology designs (Prosser et al. 2007; Knight et al. 2018), both Bray-Curtis and Aitchison distances, the latter a compositionally aware distance (Gloor et al. 2017), were used for ordination-based visualization using Principal Coordinate Analysis, linear modeling using a site-restricted, avoiding pseudo-replication, permutational analysis of variance (PERMANOVA, adonis2), and variance partitioning with partial Redundancy Analysis (RDA). Differential taxon enrichment between environmental variables was evaluated using ANCOM-BC2 (Lin and Peddada 2020), a compositionality-aware method that corrects for sampling bias and library size, with p-values adjusted for multiple testing using the Benjamini–Hochberg false discovery rate, as reviewed elsewhere (Nearing et al. 2022).

### Phylogenetic Analysis

The MAFFT software with the command: mafft --maxiterate 1000 –localpair seqs.fasta > aligned.seqs.fasta was used to generate sequence alignments (Katoh and Standley 2013). Maximum-likelihood trees with 100 bootstrap support were constructed using the RAxML program using the following parameters: raxmlHPC -f a -m GTRGAMMA -p 12345 -x 12345 -# 100 -s aligned.seqs.fasta -n T.tree, -T 4 ML search + bootstrapping (Stamatakis 2014). Newick trees files were uploaded to iTOL for visualization, labeling, and final rendering a high-quality graphics (Letunic and Bork 2021).

### Isolate Stock Generation

One cubic centimeter samples from fresh 2024 campaign mini-cores were used to generate 1:10 (wet sediment to sterile artificial seawater) slurries. One milliliter of the 1:10 slurry was aseptically spread on Zobel Marine Agar plates at sea and immediately incubated in darkness at 5°C. These incubation conditions were continued for fourteen days in the in the laboratory. We note that this approach (*i.e.*: aerobic, 5°C, in darkness) is poised to enrich psychrophilic aerobes and facultative anaerobes from the marine sediment slurry.

### Growth Kinetics and Logistic Regression

16 CFU isolates were used to generate growth curves using Zobel marine broth. Briefly, growth experiments consisted of inoculating 10 ml test tubes containing 5°C marine broth with each isolate from axenic glycerol stocks in triplicate. Test tube racks were placed in a shaker in a walk-in refrigerator (5°C) in darkness. Growth kinetics were tracked every 24 hours for 10 days measuring optical density at 600 nanometers.

The growth kinetic parameters of individual isolates were assessed to generate phenotype groupings (for example: slow vs. fast growers) and inform subsequent full-genome sequencing selection. Briefly, for each replicate, background-subtracted OD as a function of time was modeled using a three-parameter logistic growth function (Eq. 1), where K is the carrying capacity (maximum OD), r is the intrinsic growth rate, and t0 is the inflection time at which the growth rate is maximal, as detailed elsewhere (Sprouffske and Wagner 2016). Model parameters were estimated by nonlinear least squares using a Levenberg–Marquardt optimization routine, initialized with starting values based on the observed range of OD and the median time point. Model fits were inspected to ensure convergence and biological plausibility as suggested elsewhere (Sperfeld et al. 2024). To obtain a single representative growth profile per strain, parameter estimates (K, r, t0) were averaged across replicates. Distance matrices were generated from Z-standardized parameter values used as multivariate dissimilarity metrics of growth dynamics. To determine isolate with similar carrying capacity, growth rate, and timing of inflection phenotype, agglomerative hierarchical clustering was performed on the Euclidean distances using average linkage.

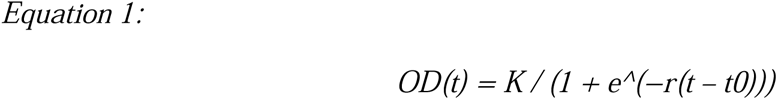

Three-parameter logistic growth model used to describe OD as a function of time, where K is the carrying capacity, r is the intrinsic growth rate, and t0 is the inflection time at which growth rate is maximal.

### Isolate Genome Sequencing and Binning

Three isolates, regrown from glycerol stocks, were sent for genome sequencing via Illumina NextSeq platform at Molecular Research DNA Labs in Lubbock, Tex. Following standard quality checks with FASTQC (Andrews 2010), SPADEs (Bankevich et al. 2012) and METABat (Kang et al. 2015) were used for contig assembly and genome binning, respectively. Genome annotations for all predicted proteins were done against the Kyoto Encyclopedia of Genes and Genomes using GhostKOALA (Kanehisa, Sato, and Morishima 2016).

### Phylogenomic Analysis and Comparative Genomics

Phylogenomic analysis of isolate genomes was performed using the GToTree package (Lee 2019). Briefly, our subseafloor isolate genomes and 4,852 Gammaproteobacterial public genomes, randomly selected from 226,629 Gammaproteobacteria genome entries available in the Genome Taxonomy Database (GTDB v226) released Apr 16, 2025, were analyzed with GToTree’s “Gammaproteobacteria” profile Hidden Markov Model (pHMM) collection of lineage-specific single copy genes. Identified homologues were aligned using Muscle (Edgar 2004) resulting in a concatenated protein alignment subsequently trimmed with TrimAl (Capella-Gutiérrez, Silla-Martínez, and Gabaldón 2009). FastTree2 (Price, Dehal, and Arkin 2010) was used for tree construction and visualizations were performed on iTOL (Letunic and Bork 2021). Subseafloor isolates and closest relative genomes were compared using predicted KEGG Decoder’s metabolic pathway prediction summaries (Graham, Heidelberg, and Tully 2018). Biosynthetic gene clusters for secondary metabolite predictions were identified with antiSMASHv6.0 (Blin et al. 2021).

## Results and Discussion

### Oceanographic Setting and Interpretative Context

A total of six sediment cores were recovered across the San Pedro channel slopes during the 2023 and 2024 coring campaigns (Figure 1). Cross-channel coring occurred at standardized water depths and, when possible, mini-core subsampling and sequencing was also standardized to depth beneath the seafloor, as discussed below. Despite subseafloor and water depth standardization, differences in sediment history in across spatial scales prohibits direct across-slope ecological comparisons. Differences in sedimentation rate (Covault et al. 2007), basin topography (Normark et al. 2009), and organic matter source (Romans et al. 2009) result in natural environmental variability at the pelagic-benthic interface and ultimately establish ecological boundaries in subseafloor sediment (D’Hondt, Rutherford, and Spivack 2002; D’Hondt et al. 2019). This complexity is increasingly recognized as essential to understanding how microbial communities respond to coupled physical, chemical, and anthropogenic influences in subseafloor environments at regional scales (Orcutt et al. 2011).

Cross-basin slope boundary comparisons beneath one of the world most impacted maritime corridors represent a natural experiment where, despite environmental heterogeneity and the limited number of samples proffered by this study, a subseafloor anthropogenic signature may be eventually explored. This regional coring is an initial effort to characterize subseafloor microbiological baselines under different depositional (near-shore vs. mid-channel) and human disturbance (Los Angeles County vs. Santa Catalina Island slopes) regimes in the San Pedro basin.

### Regional Subseafloor Ecology Survey: Community Diversity

As expected in this environmentally complex sample collection, the Shannon-Weaver alpha diversity index varies systematically between slopes and depth beneath the seafloor (Figure 2A). The reported values match equivalent siliciclastic clay horizons collected between Santa Catalina and San Clemente Islands, in the Borderlands region (Ramírez, Graham, and D’Hondt 2018). Despite a general drop in alpha diversity with depth beneath the seafloor (bsf) within each slope at each water depth, a differential trend emerges: at most depth bsf matched pairings, samples of the San Pedro slope showed a more pronounced reduction in diversity and/or evenness relative to equivalent depths bsf in the Santa Catalina Island slope, particularly in shallow-water depth sites (Figure 2A, 65 m). Midwater samples display depth bsf and variance-related distinctions in Shannon-Weaver index values across slopes: Santa Catalina slope communities show a broader alpha diversity index spread, whereas this index is more clustered for San Pedro communities (Figure 2A, 401 m). At the channel bottom, two out of three depth bsf pairs show Santa Catalina communities retaining higher alpha diversity index values (Figure 2A, 893 m).

**Figure 2.**
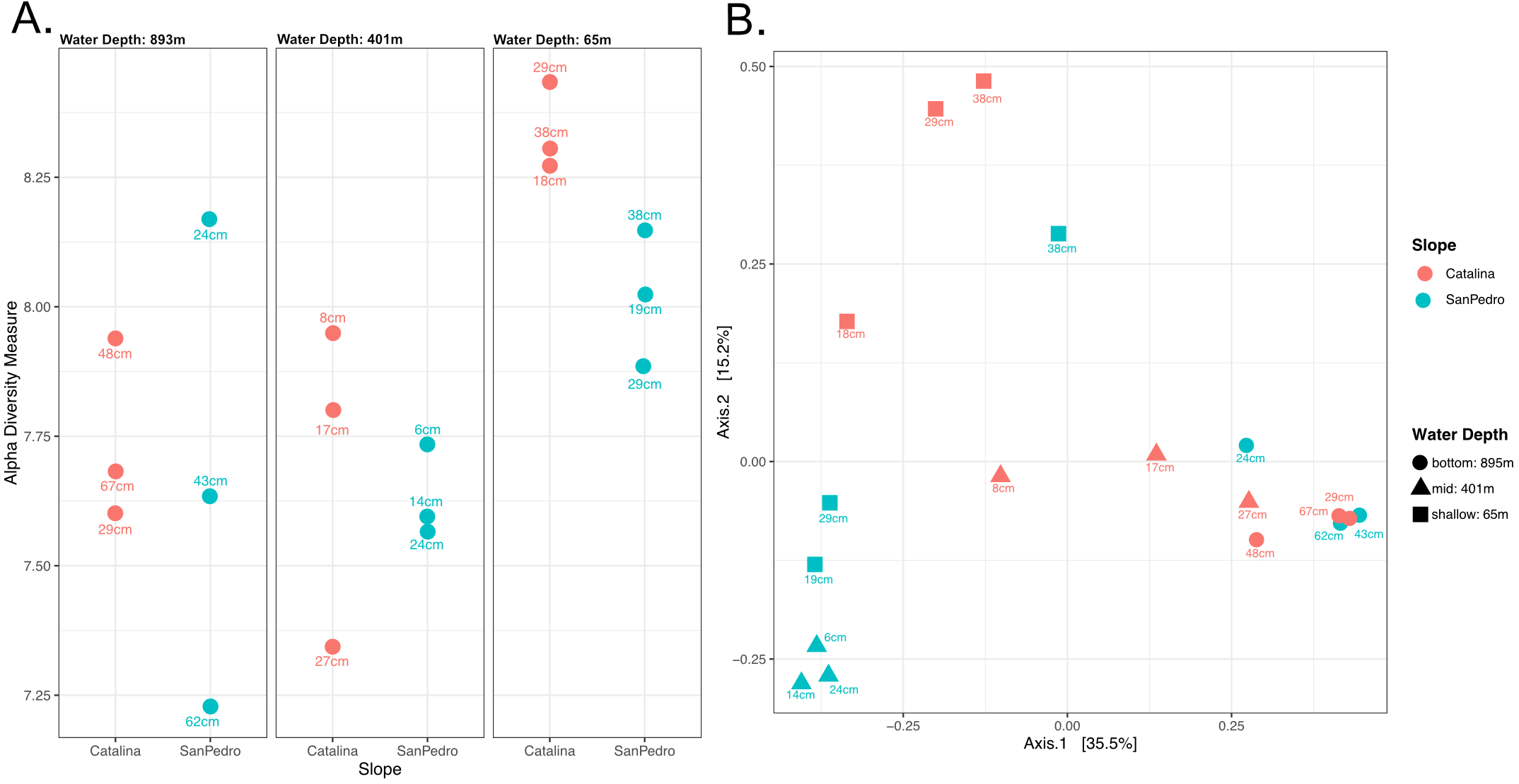
Diversity metrics. A) Shannon alpha diversity. Each panel shows sample data for each water depth, with color depicting Santa Catalina (red) or San Pedro (teal) slope provenance and shape depicting water depth at each sampled site. Depth in cm beneath the seafloor (cm bsf) shown in icon labels. B) Ordination (PCoA) summary displayed with identical color, shape, and label descriptors as panel A.

Focusing on emergent structural patterns, ordination visualization suggests water separation across axis 1, explaining 35.5% of the variance, while axis 2, explaining 15.2 % of the variance, separates samples generally based on Santa Catalina Island versus San Pedro basin slopes (Figure 2B). Permutational Analysis of Variance shows overlaying water depth, independently of site, as the strongest determinant of community composition (Table 1). Overall, these ecological predictors are concordantly significant when analyzed with abundance-weighted Bray-Curtis and the compositionally-aware Aitchison distances with close rank order of effect sizes strongly suggesting that our observations are not driven by compositional artifacts.

**Table 1.** Comparison of PERMANOVA results using Bray-Curtis and Aitchison distances.

| Predictor | Bray-Curtis $R^2$ | Aitchison $R^2$ | Bray-Curtis p-value | Aitchison p-value | Stability across metrics |
| --- | --- | --- | --- | --- | --- |
| Slope | 0.085 | 0.080 | 0.001 | 0.001 | Highly Similar |
| Water Depth | 0.185 | 0.159 | 0.001 | 0.001 | Highly Similar |
| Depth Beneath Seafloor | 0.091 | 0.081 | 0.025 | 0.050 | Persistent |
| Residual | 0.547 | 0.627 | - | - | Unexplained Variance |

Previous surficial sediment work in the region has shown that despite stable community abundance and diversity metrics observed across environmental and geographical scales, the composition of pelagic-benthic interface (>1 cm bsf) communities may change drastically across short spatial scales (Hewson, Jacobson Meyers, and Fuhrman 2007). Here, we report both structural and compositional differences as a function of environmental gradients in the San Pedro Basin subseafloor (Figure 2). Specifically, our results, which target significantly deeper sediment depth horizons (6-67 cm bsf) relative to the previous work (>1 cm bsf), suggest that water depth is the strongest driver of subseafloor community assembly in the San Pedro Basin (Figure 2 & Table 1). Others have identified large-scale ecological patterns in pelagic microbial communities (DeLong et al. 2006), with convergent diversity peaks at redox-defined oceanographic features, such as oxygen minimum zones, across ocean basins (Walsh et al. 2016). Accordingly, water depth, likely through depth- and seasonality-dependent structuring of the regional water column microbial community (Cram et al. 2015), determines the pool of lineages available for seeding the seafloor, thereby influencing subseafloor microbial community structure in the San Pedro Basin region. This mechanism explains both the widespread structural similarity of sedimentary communities at the pelagic-benthic interface (Hewson, Jacobson Meyers, and Fuhrman 2007) and the environment-specific (water depth) differentiation in subseafloor microbial assemblages after decades to centuries of burial (Figure 2).

Interestingly, at equivalent water and subseafloor depths, slope plays a significant role in microbial community structuring (Figure 2A & Table 1). Because our sampling design omits environmental factors (*e.g.* geochemistry, grain size, mineralogy) that may also exert significant ecological effects (Warrick and Farnsworth 2009), slope-driven differences are only interpreted as a persistent regional effect on subseafloor microbial communities. Subsurface depth, ultimately reflecting biogeochemical microbial respiration niches, has a secondary but significant effect on subseafloor community structure in the region (Figure 2B & Table 1). We note that, due to a lower attributable fraction of the variance under the Aitchison framework (Table 1), depth beneath seafloor appears to primarily influence relative abundances rather than taxonomic turnover in this survey.

### Regional Subseafloor Ecology Survey: Community Composition

Dominant phyla across all samples include Acidobacteria, Asgardarchaeota, Atribacterota, Chloroflexota, Planctomycetota, Pseudomonadota, Thermodesulfobacteriota, Thermoplasmatota, and Thermoproteota with a Bacteria to Archaea geometric mean of 3.22 (Figure 3). These are classical subseafloor lineages generally associated with continental slope sedimentary habitats (Orcutt et al. 2011; Orsi 2018), closely matching microbial community structure from similar depth horizons from elsewhere in the Borderlands region analyzed using identical extraction, amplification, and sequencing techniques (Ramírez et al. 2018). Shallow (65 m water depth) communities display relatively balanced phylum level representation, consistent with alpha diversity estimates. As expected from ordination visualizations (Figure 2B), Midwater (405 m water depth) samples show more pronounced variation across slopes, driven by differences in the relative abundance of groups such as Thermoproteota, Thermodesulfobacteriota, and Bacteroidota, while channel bottom (893 m water depth) communities were the most compositionally similar across the transect (Figure 3).

**Figure 3.**
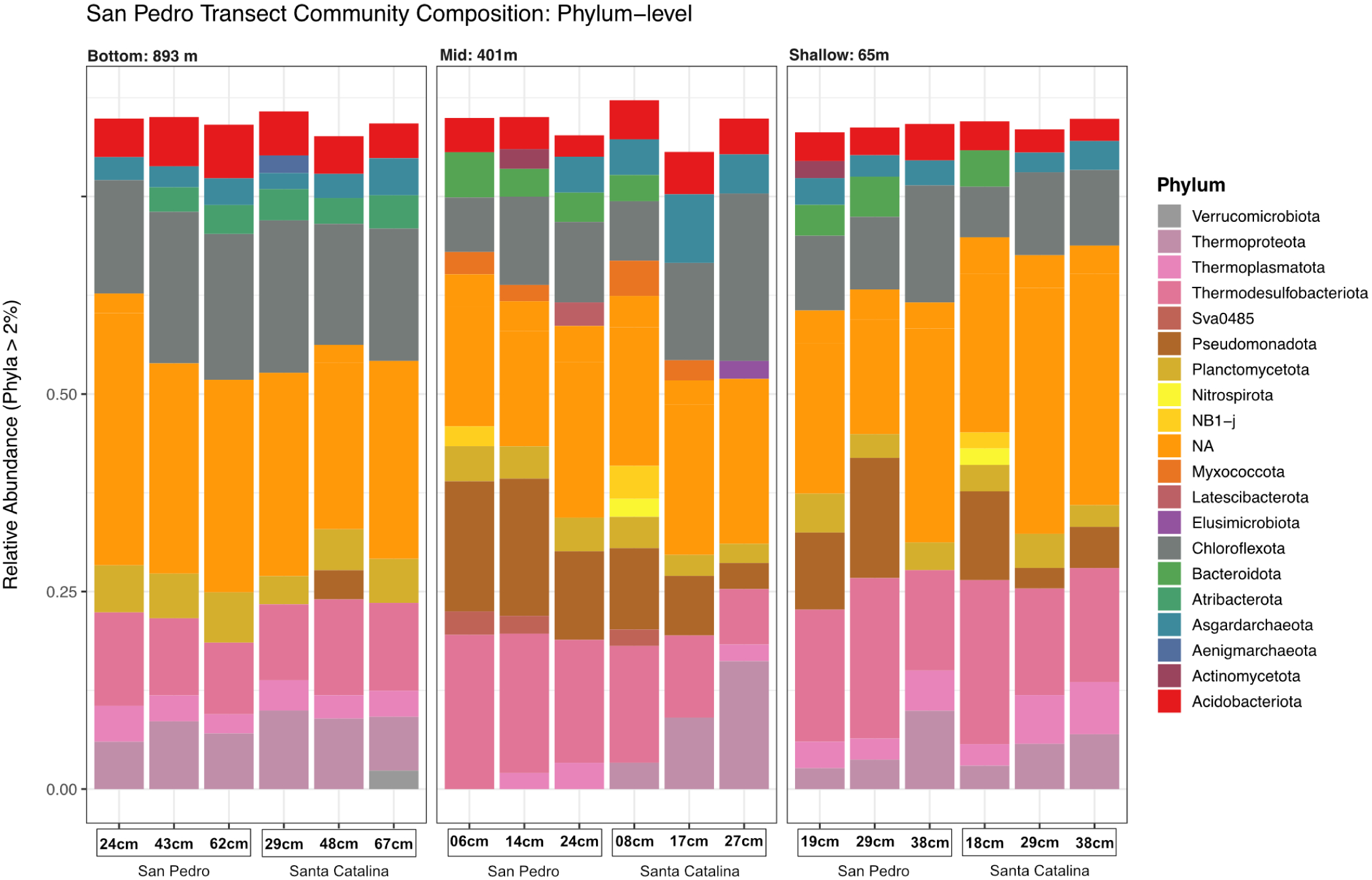
Phylum level Microbial Community Composition. The left, middle, right depict sedimentary microbial communities collected from 893 m, 401 m, and 65 m water depth environments. Within each panel, column labels show depth beneath the seafloor in centimeters and either San Pedro or Santa Catalina slope collection site.

Based on our previous work in the area, all samples in this survey are expected to be anaerobic environments with high organic matter content [2-4% Total Organic Carbon (TOC)] with nitrate oxidation in relatively shallow subseafloor depths (Ramírez et al. 2018). Collectively, our survey supports anaerobic heterotrophy and fermentation coupled with sulfate reduction under high TOC but rather energy-limited conditions (Figure 3). The Chloroflexota, widely abundant in our survey, are a bacterial heterotrophic lineage characteristic of low energy subseafloor habitats capable of syntrophic interactions and hydrogen cycling, well-adapted to low energy requirements (Blazejak and Schippers 2010). High abundance members of this lineage, belonging to the classes Dehalococcoidia and Anaerolineae (Figure 4), notorious for organohalide respiration (Yang et al. 2020), are close relatives of clones reported from a sulfate-methane transition zone (SMTZ) study in the nearby Santa Barbara Basin (Harrison et al. 2009) and the Sonora Margin, Gulf of California (Vigneron et al. 2013). Other dominant ASVs were identified as members of the Thermodesulfobacteriota class Syntrophobacteria, obligate anaerobes generally coupled with hydrogenotrophic methanogens and/or sulfate reducing bacteria (Waite et al. 2020), were closely related to sequences recovered from a Nankai Trough methanogenic sediment enrichment (Aoki et al. 2014) and off-shore Shimokita drill cores (Kawai et al. 2014) from the Western Pacific (Figure 4). Members of the Desulfobacterota phylum, comprised of strictly anaerobic sulfate reducers (Waite et al. 2020), were dominant across all samples in this survey (Figures 3 & 4). Abundant members of this phylum include ASVs classified as Desulfobacteria (Waite et al. 2020), strict sulfate reducers capable or complete organic carbon remineralization, closely related to clones recovered from globally-distributed sites including the Western Pacific (Kawai et al. 2014), Indian Ocean, and Gulf of México (Nunoura et al. 2009). Another member of this phylum, belonging to the class Desulfobulbia (Waite et al. 2020), is a close relative of a sequence recovered from Arctic sediment (Bowman and McCuaig 2003). Other high abundance ASVs included members of the Atribacterota (Class JS1) and Aminicenantia (formally OP8) (Figures 3 & 4), anaerobes driving carbon and nitrogen, via fermentations and necromass turnover, over long timescales.

**Figure 4.**
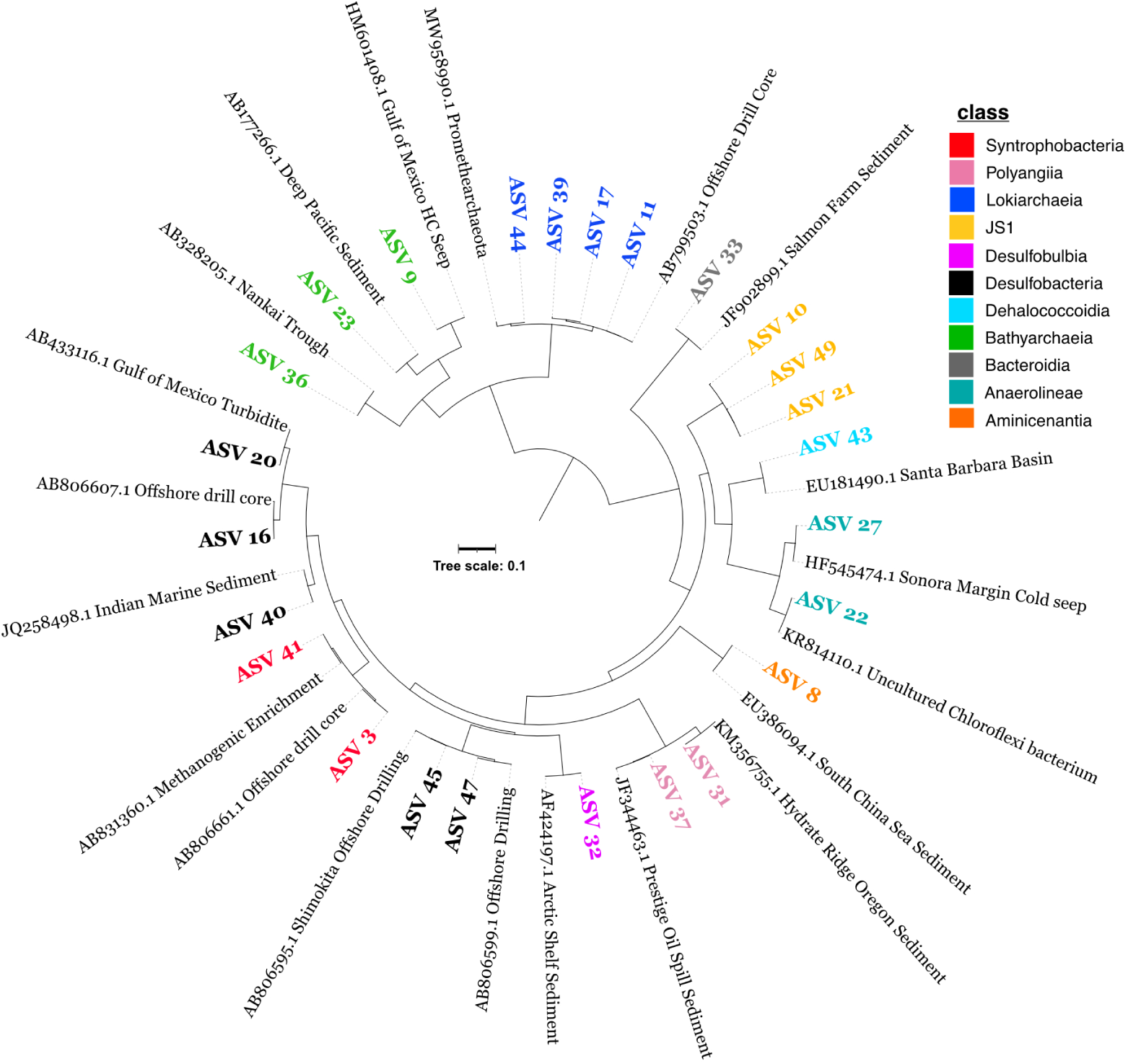
Phylogenetic analysis of high abundance Amplicon Sequence Variants (ASVs). Colors depict the taxonomic affiliation of each ASV. Neighboring leaves represent closest geographically annotated relatives based on BLASTn results generated in November 2025.

Archaea were also abundant in this survey (Figure 3). Dominant lineages included members of the Lokiarchaeia and Bathyarchaeia classes (Figure 4). Lokiarchaeia ASVs are close relatives of the first culture representative of this lineage “*Candidatus* Prometheoarchaeum syntrophicum” strain MK-D1 (Avci et al. 2022). Representative genomes suggest that the Lokiarchaeia is comprised of sedimentary heterotrophs capable of degrading complex organic matter in anoxic sediment (Yin et al. 2021; Spang et al. 2015). Bathyarchaeia ASVs are related to clone sequences from methane hydrate-bearing Pacific deep sediment (Inagaki et al. 2006) and seeps from the Gulf of México (Lanoil et al. 2001). Bathyarchaeia [former Miscellaneous Crenarchaeotal group (MCG)] are widespread in marine sediment and contain representatives capable of detrital protein degradation (Lloyd et al. 2013). Together, these lineages, are involved in carbon and nitrogen cycling in the San Pedro Basin subseafloor.

### Physiological Characterization of Subseafloor Isolates

The subseafloor, comprised of vertically stratified redox niches, harbors a significant portion of the planet’s biodiversity (Hoshino et al. 2020). Unsurprisingly, marine sediment is rapidly emerging as a potential treasure trove of novel natural products with biotechnological applications (Bech et al. 2020; Orsi, Richards, and Francis 2018). Targeted cultivation of subseafloor isolates, however, is not trivial. Habitat complexity, including adaption to extreme energy-limitation, selects for denizens with fastidious phenotypes, necessitating sophisticated culturing setups and/or long incubation times for successful targeted microbiological enrichment and isolation (Imachi et al. 2020). Despite this, the vast number and diversity of cells in marine sediment also provide a powerful opportunity for isolate recovery. Leveraging on this fact, untargeted cultivation approaches can yield diverse and phylogenetically novel isolates amendable to sequencing and genomic characterized (Russell et al. 2016; Orsi, Magritsch, Vargas, Coskun, Vuillemin, Höhna, Wörheide, D’Hondt, et al. 2021) but, due to extremely slow growth rates (Orsi 2018; Hoehler and Jørgensen 2013), such efforts are rather rare.

We successfully recovered hundreds of colony forming units (CFUs) from subseafloor sediment slurries plated on marine media in aerobic 5 °C conditions in darkness (data not shown). Fifteen of these CFUs, based on unique colony morphology, were selected for growth analysis. Physiological characterization of our subseafloor isolates revealed substantial heterogeneity in growth dynamics under low-temperature (5 °C) and aerobic conditions. Some lineages show rapid early growth, some exhibit prolonged lag phases, and others display little to no growth (Figure 5A). These measured growth phenotypes display the physiological diversity in rate, carrying capacity, and timing of maximal growth of subseafloor denizens. Clustering based on logistic growth parameters (K, r, and t0) resolved isolates into distinct growth phenotypes (Figure 5B). Isolates group based on fast growing/high yield versus slow growth low yield, both potential adaptive or selected responses to high-carbon/low-energy habitats (Vuillemin et al. 2020). Notably, isolates selected for whole-genome sequencing represent different growth clusters (Figure 5B, denoted by a star) capturing both relatively active and growth-limited phenotypes. Together, these results demonstrate that subseafloor isolates retain diverse physiological strategies consistent with niche differentiation and variable energetic constraints and provide an initial catalogue of growth-characterized models from the San Pedro Basin subseafloor for future experimental work.

**Figure 5.**
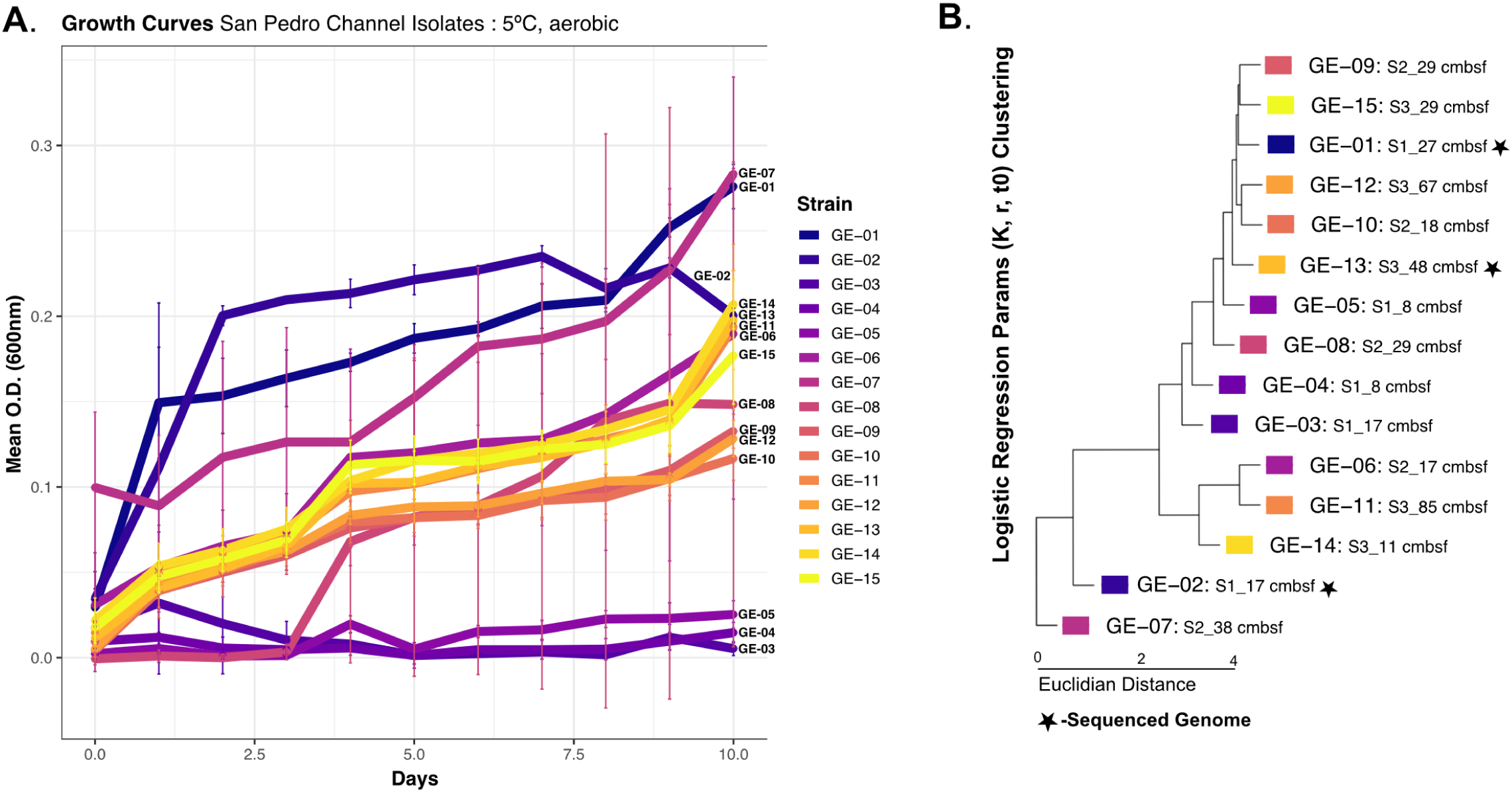
Isolate Growth Kinetics. A) Growth curve data for 15 subseafloor isolates. B) Growth phenotype (K, r, t_0_) clustering for the 15 isolate strains. Tree leaves depict isolate ID (GE-x): 2024 collection site (Sx_) and depth beneath seafloor (x cmbsf), with whole-genome sequencing selections denoted by star symbols.

### Comparative Genomics of Subseafloor Isolates

Sequenced isolates from our subseafloor microbiological collection were classified as Gammaproteobacteria. To place these genomes in an evolutionary context, we generated a large phylogenetic tree using 4,852 Gammaproteobacterial genomes from GTDB (Figure 6A). This analysis restricted our isolate placement to a relatively narrow region of the large Gammaproteobacteria tree. The recovery of closely-related Gammaproteobacteria lineages reflects well-known methodological (incubation conditions and media type) bias which preferentially select for heterotrophic lineages (Bech et al. 2020) rather than the full breadth of the natural sediment community (Figure 3).

**Figure 6.**
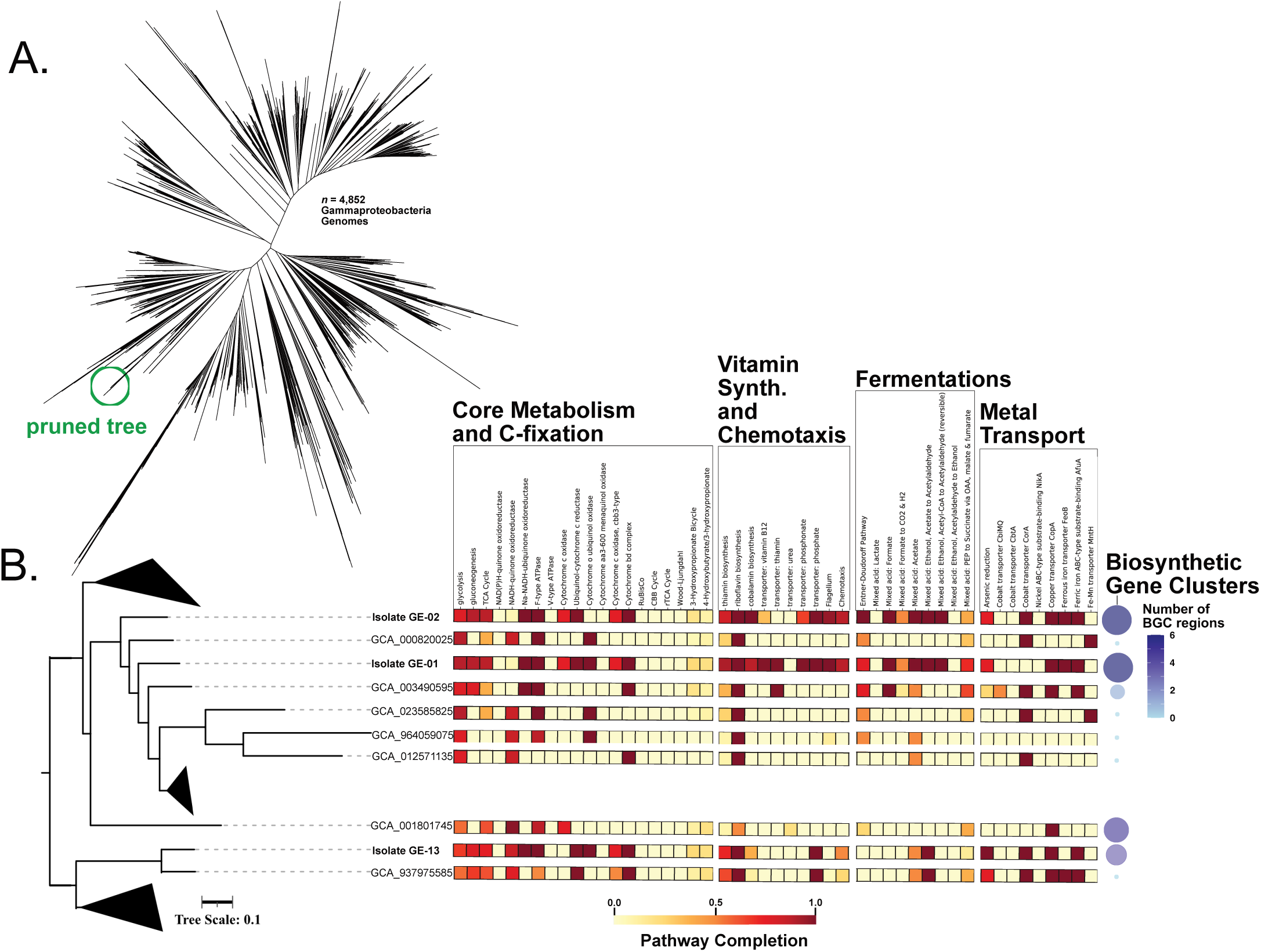
Phylogenomic and comparative genomic analysis. A) Unrooted phylogenomic tree of Gammaproteobacterial high quality and completion (>98%) genomes generated from our isolate collection and 4,852 randomly selected lineages from GTDB. B) Pruned phylogenomic tree with functional predictions for core metabolism, carbon fixation, vitamin synthesis, chemotaxis, fermentations, and metal transport depicted as a yellow to red color scale pathway completion heatmap. Predicted Biosynthetic Gene Clusters for each genome are summarized in the blue-scale right-most dot plot column.

Focusing on the closest-related reference genomes show that the isolates share a common core metabolic repertoire, including pathways for central carbon metabolism, predicted heterotrophy (no carbon fixation), and the biosynthesis of essential B-vitamins (thiamin, riboflavin, and cobalamin), while differing in traits such as chemotaxis, fermentation pathways, and metal transport systems (Figure 6B). Differences in isolate genomes likely reflect niche-specific adaptations to variable redox conditions, substrate availability, and trace metal concentrations in subseafloor sediments, as suggested elsewhere (Orsi 2018; Orsi, Richards, and Francis 2018). Interestingly, isolates GE-01 and -02, recovered from 65m water depth and 27 and 17 cm bsf, respectively, retain flagellar biosynthesis genes, highlighting their pelagic provenance. In contrasts, isolate genome GE-13, recovered from full-channel water depth and approximately half a meter bsf, does not have flagellar biosynthesis genes (Fig. 6B), a previously reported ecological strategy expected for microorganisms persisting long-term under diffusion-limited energy limitation (Hoehler and Jørgensen 2013).

We report that subseafloor isolate genomes contained significantly (student t-test; Pval<0.05) more biosynthetic gene clusters (BGCs) compared to their closest analyzed relatives from GTDB (Figure 6B, Table 2). Predicted BGCs from subseafloor include the following product classes: Nonribosomal Peptide Synthetase (NRPS), Ribosomally synthesized and post-translationally modified peptides (RiPP), Arylpolyene, Polyunsaturated fatty acids (PUFA), siderophores, glycolipid-like ketosynthase (hglE-KS) and, most commonly, beta-lactone biosynthetic clusters (Table 2). PUFA and hglE-KS predicted products suggests membrane modification leading to fluidity modification and/or surface adhesion as an important adaptive theme subseafloor genomes (Lever et al. 2015). Predictions of siderophores (Garber et al. 2021), lipid-linked arylpolyene pigments (Dong et al. 2024), and RiPP-like gene clusters suggests iron-acquisition, oxidative stress protection, and redox balancing, as subseafloor activities by these lineages. Lastly, beta-lactones and NRPS clusters strongly suggest microbe-microbe interactions particularly biocidal product-based antagonistic competition, all important expected activities in diffusion-limited, low-energy subseafloor sediment (Hoehler and Jørgensen 2013; Lever et al. 2015).

**Table 2.** Biosynthetic Gene Cluster (BGC) counts and description for genomes from. **Figure 6B**.

| Genome | # BGC | # BGC Classes | BGC Class Description |
| --- | --- | --- | --- |
| GCA_003490595 | 1 | 1 | LAP (RiPP) |
| GCA_012571135 | 0 | 0 | none detected |
| GCA_001801745 | 4 | 4 | NRPS-like; RiPP-like; terpene; arylpolyene |
| GCA_937975585 | 0 | 0 | none detected |
| GCA_023585825 | 0 | 0 | none detected |
| GCA_964059075 | 0 | 0 | none detected |
| GE-1 (GAR19) | 6 | 6 | betalactone; NRPS; arylpolyene; siderophore |
| GE-2 (GAR20) | 6 | 6 | RiPP-like; arylpolyene; betalactone; hglE-KS; PUFA |
| GE-13 (GAR21) | 3 | 3 | siderophore; redox-cofactor (RiPP); betalactone |

## Conclusions

Microbial communities inhabiting marine sediments represent a substantial portion of Earth’s biomass and are central to global biogeochemical processes. Yet, the subseafloor ecosystems of the San Pedro Channel—situated between Los Angeles and Santa Catalina Island—remain largely unexplored, despite the region’s long history of intense maritime activity. To begin addressing this knowledge gap, we initiated an oceanographic sediment coring program that produced a depth standardized transect across both the Los Angeles and Catalina slopes. Rather than serving as a direct comparison of the basin margins, this effort establishes an ecological reference point for the San Pedro Basin subseafloor that can support future regional environmental monitoring efforts and provides a curated collection of representative subseafloor isolates for experimental and comparative genomic analyses.

## Acknowledgements

We thank the crew of the Southern California Marine Institute Research Vessel (R/V) Yellowfin for technical assistance during coring operations. We also thank the California State University, Los Angeles, Applied and Environmental Microbiology undergraduates that sailed in the 2023-2024 legs as student observers and assisted with at-sea core handling and microbiological operations.

## Works Cited

Ahn, Jong Ho, and Stanley B. Grant. 2007. ‘Characteristics of storm runoff and sediment dispersal in the San Pedro Channel, southern California’, Water Science and Technology, 55: 519–26.

Andrews, S. 2010. ‘FastQC: a quality control tool for high throughput sequence data.’.

Aoki, Masataka, Masayuki Ehara, Yumi Saito, Hideyoshi Yoshioka, Masayuki Miyazaki, Yayoi Saito, Ai Miyashita, Shuji Kawakami, Takashi Yamaguchi, Akiyoshi Ohashi, Takuro Nunoura, Ken Takai, and Hiroyuki Imachi. 2014. ‘A Long-Term Cultivation of an Anaerobic Methane-Oxidizing Microbial Community from Deep-Sea Methane-Seep Sediment Using a Continuous-Flow Bioreactor’, PLoS One, 9: e105356.

Avci, Burak, Jakob Brandt, Dikla Nachmias, Natalie Elia, Mads Albertsen, Thijs J. G. Ettema, Andreas Schramm, and Kasper Urup Kjeldsen. 2022. ‘Spatial separation of ribosomes and DNA in Asgard archaeal cells’, The ISME Journal, 16: 606–10.

Bankevich, Anton, Sergey Nurk, Dmitry Antipov, Alexey A. Gurevich, Mikhail Dvorkin, Alexander S. Kulikov, Valery M. Lesin, Sergey I. Nikolenko, Son Pham, Andrey D. Prjibelski, Alexey V. Pyshkin, Alexander V. Sirotkin, Nikolay Vyahhi, Glenn Tesler, Max A. Alekseyev, and Pavel A. Pevzner. 2012. ‘SPAdes: A new genome assembly algorithm and its applications to single-cell sequencing’, Journal of Computational Biology, 19: 455–77.

Bar-On, Yinon M., Rob Phillips, and Ron Milo. 2018. ‘The biomass distribution on Earth’, Proceedings of the National Academy of Sciences of the United States of America, 115: 6506–11.

Bech, Pernille Kjersgaard, Klaus Lars Lysdal, Lone Gram, Mikkel Bentzon-Tilia, and Mikael Lenz Strube. 2020. ‘Marine Sediments Hold an Untapped Potential for Novel Taxonomic and Bioactive Bacterial Diversity’, mSystems, 5: e00782–20.

Blazejak, Anna, and Axel Schippers. 2010. ‘High abundance of JS-1- and Chloroflexi-related bacteria in deeply buried marine sediments revealed by quantitative, real-time PCR’, FEMS Microbiology Ecology, 72: 198–207.

Blin, Kai, Simon Shaw, Anne M. Kloosterman, Zachary Charlop-Powers, Gilles P. van Wezel, Marnix H. Medema, and Tilmann Weber. 2021. ‘antiSMASH 6.0: improving cluster detection and comparison capabilities’, Nucleic Acids Research, 49: W29–W35.

Bowman, John P., and Robert D. McCuaig. 2003. ‘Biodiversity, community structural shifts, and biogeography of prokaryotes within Antarctic continental shelf sediment’, Applied and Environmental Microbiology, 69: 2463–83.

Bradley, J.□A., J.□P. Amend, D.□E. LaRowe, S. Ono, S. D’Hondt, and B.□N. Orcutt. 2020. ‘Widespread energy limitation to life in global subseafloor sediments’, Science Advances, 6: eaba0697.

California Regional Water Quality Control Board, Los Angeles Region. 2016. “Order R4-2016-0334: Waste Discharge Requirements and NPDES Permit for Terminal Island Water Reclamation Plant (TIWRP).” In.: State Water Resources Control Board, California.

Callahan, B. J., P. J. McMurdie, and S. P. Holmes. 2017. ‘Exact sequence variants should replace operational taxonomic units in marker-gene data analysis’, ISME Journal, 11: 2639–43.

Callahan, B. J., P. J. McMurdie, M. J. Rosen, A. W. Han, A. J. Johnson, and S. P. Holmes. 2016. ‘DADA2: High-resolution sample inference from Illumina amplicon data’, Nat Methods, 13: 581–3.

Capella-Gutiérrez, Salvador, José M. Silla-Martínez, and Toni Gabaldón. 2009. ‘trimAl: a tool for automated alignment trimming in large-scale phylogenetic analyses’, Bioinformatics, 25: 1972–73.

Covault, J. A., W. R. Normark, B. W. Romans, and S. A. Graham. 2007. ‘Highstand fans in the California Borderland: The overlooked deep-water depositional systems’, Geology, 35: 783–86.

Cram, Jacob A., Cheryl-Emiliane T. Chow, Rohan Sachdeva, David M. Needham, Alma E. Parada, Joshua A. Steele, and Jed A. Fuhrman. 2015. ‘Seasonal and interannual variability of the marine bacterioplankton community throughout the water column over ten years’, The ISME Journal, 9: 563–80.

D’Hondt, S., B.B. Jørgensen, D.J. Miller, A. Batzke, R. Blake, B.A. Cragg, H. Cypionka, G.R. Dickens, T. Ferdelman, K.-U. Hinrichs, N.G. Holm, R. Mitterer, A. Spivack, G. Wang, B. Bekins, B. Engelen, K. Ford, G. Gettemy, S.D. Rutherford, H. Sass, C.G. Skilbeck, I.W. Aiello, G. Guèrin, C.H. House, F. Inagaki, P. Meister, T. Naehr, S. Niitsuma, R.J. Parkes, A. Schippers, D.C. Smith, A. Teske, J. Wiegel, C.N. Padilla, and J.L.S. Acosta. 2004. ‘Distributions of microbial activities in deep subseafloor sediments’, Science, 306: 2216–21.

D’Hondt, S., S. Rutherford, and A. J. Spivack. 2002. ‘Metabolic activity of subsurface life in deep-sea sediments’, Science, 295: 2067–70.

D’Hondt, Steven, Robert Pockalny, Victoria M. Fulfer, and Arthur J. Spivack. 2019. ‘Subseafloor life and its biogeochemical impacts’, Nature Communications, 10.

Davis, N.□M., D.□M. Proctor, S.□P. Holmes, D.□A. Relman, and B.□J. Callahan. 2018. ‘Simple statistical identification and removal of contaminant sequences in marker□gene and metagenomics data’, Microbiome, 6: 226.

DeLong, Edward F., Christina M. Preston, Tracy Mincer, Virginia Rich, Steven J. Hallam, Niels Ulrik Frigaard, Asuncion Martinez, Matthew B. Sullivan, Robert Edwards, Beltran Rodriguez Brito, Sallie W. Chisholm, and David M. Karl. 2006. ‘Community genomics among stratified microbial assemblages in the ocean’s interior’, Science, 311: 496–503.

Dong, Xiyang, Tianxueyu Zhang, Weichao Wu, Yongyi Peng, Xinyue Liu, Yingchun Han, Xiangwei Chen, Zhizeng Gao, Jinmei Xia, Zongze Shao, and Chris Greening. 2024. ‘A vast repertoire of secondary metabolites potentially influences community dynamics and biogeochemical processes in cold seeps’, Science Advances, 10: eadl2281.

Edgar, Robert C. 2004. ‘MUSCLE: multiple sequence alignment with high accuracy and high throughput’, Nucleic Acids Research, 32: 1792–97.

Eganhouse, Robert P., and John Pontolillo. 2000. ‘Depositional history of organic contaminants on the Palos Verdes Shelf, California’, Marine Chemistry, 70: 317–38.

Fisher, Michael A., William R. Normark, Victoria E. Langenheim, Andrew J. Calvert, and Ray W. Sliter. 2004. “Marine geology and earthquake hazards of the San Pedro Shelf region, Southern California.” In U.S. Geological Survey Professional Paper. Reston, VA: U.S. Geological Survey.

Garber, A.□I., A.□B. Cohen, K.□H. Nealson, G.□A. Ramírez, R.□A. Barco, T.□C. Enzingmüller Bleyl, M.□M. Gehringer, and N. Merino. 2021. ‘Metagenomic Insights Into the Microbial Iron Cycle of Subseafloor Habitats’, Frontiers in Microbiology, 12: 667944.

Garrett, Chris, and Eric Kunze. 2007. ‘Internal tide generation in the deep ocean’, Annual Review of Fluid Mechanics, 39: 57–87.

Gloor, Gregory B., Jean M. Macklaim, Vera Pawlowsky-Glahn, Juan J. Egozcue, Thomas P. Quinn, and Andrew D. Fernandes. 2017. ‘Microbiome Datasets Are Compositional: And This Is Not Optional’, Frontiers in Microbiology, 8: 2224.

Graham, Emily D., John F. Heidelberg, and Benjamin J. Tully. 2018. ‘Potential for primary productivity in a globally-distributed bacterial phototroph’, ISME Journal, 12: 1861–66.

Harrison, B. K., H. Zhang, W. Berelson, and V. J. Orphan. 2009. ‘Variations in archaeal and bacterial diversity associated with the sulfate-methane transition zone in continental margin sediments (Santa Barbara Basin, California)’, Applied and Environmental Microbiology, 75: 1487–99.

Hewson, Ian, Marcia E. Jacobson Meyers, and Jed A. Fuhrman. 2007. ‘Diversity and biogeography of bacterial assemblages in surface sediments across the San Pedro Basin, Southern California Borderlands’, Environmental Microbiology, 9: 923–33.

Hoehler, T.□M., and B.□B. Jørgensen. 2013. ‘Microbial life under extreme energy limitation’, Nature Reviews Microbiology, 11: 83–94.

Hoshino, Takuya, Hideyuki Doi, Guiomar I. Uramoto, Liane Wörmer, Yuki Matsui, Kazuhiro Hori, Masayuki Miyazaki, and, et al. 2020. ‘Global diversity of microbial communities in marine sediment’, Proceedings of the National Academy of Sciences of the United States of America, 117: 27587–97.

Hoshino, Tatsuhiko, and Fumio Inagaki. 2019. ‘Abundance and distribution of Archaea in the subseafloor sedimentary biosphere’, The ISME Journal, 13: 227–31.

Imachi, Hiroyuki, Masaru K. Nobu, Nozomi Nakahara, Yuki Morono, Miyuki Ogawara, Yoshihiro Takaki, Yoshinori Takano, Katsuyuki Uematsu, Tetsuro Ikuta, Motoo Ito, Yohei Matsui, Masayuki Miyazaki, Kazuyoshi Murata, Yumi Saito, Sanae Sakai, Chihong Song, Eiji Tasumi, Yuko Yamanaka, Takashi Yamaguchi, Yoichi Kamagata, Hideyuki Tamaki, and Ken Takai. 2020. ‘Isolation of an archaeon at the prokaryote–eukaryote interface’, Nature, 577: 519–25.

Inagaki, Fumio, Mitsuo Suzuki, Ken Takai, Hiroshi Oida, Toshiaki Sakamoto, and Kazuo Aoki. 2006. ‘Biogeographical distribution and diversity of microbes in methane hydrate-bearing deep marine sediments on the Pacific Ocean Margin’, Proceedings of the National Academy of Sciences of the United States of America, 103: 2815–20.

Jorgensen, B. B., and I. P. G. Marshall. 2016. ‘Slow microbial life in the seabed’, Annual Review of Marine Science, 8: 311–32.

Kallmeyer, Jens, Robert Pockalny, Rishi R. Adhikari, David C. Smith, and Steven D’Hondt. 2012. ‘Global distribution of microbial abundance and biomass in subseafloor sediment’, Proceedings of the National Academy of Sciences of the United States of America, 109: 16213–16.

Kanehisa, Minoru, Yoko Sato, and Kanako Morishima. 2016. ‘BlastKOALA and GhostKOALA: KEGG tools for functional characterization of genome and metagenome sequences’, Journal of Molecular Biology, 428: 726–31.

Kang, Dongwan D., Jeff Froula, Rob Egan, and Zhong Wang. 2015. ‘MetaBAT, an efficient tool for accurately reconstructing single genomes from complex microbial communities’, PeerJ, 3: e1165.

Katoh, Kazutaka, and Daron M. Standley. 2013. ‘MAFFT multiple sequence alignment software version 7: improvements in performance and usability’, Molecular Biology and Evolution, 30: 772–80.

Kawai, Mikihiko, Taiki Futagami, Atsushi Toyoda, Yoshihiro Takaki, Shinro Nishi, Sayaka Hori, Wataru Arai, Taishi Tsubouchi, Yuki Morono, Ikuo Uchiyama, Takehiko Ito, Asao Fujiyama, Fumio Inagaki, and Hideto Takami. 2014. ‘High frequency of phylogenetically diverse reductive dehalogenase-homologous genes in deep subseafloor sedimentary metagenomes’, Frontiers in Microbiology, 5: 80.

Kirkpatrick, J. B., E. A. Walsh, and S. D’Hondt. 2019. ‘Microbial Selection and Survival in Subseafloor Sediment’, Frontiers in Microbiology, 10: 956.

Knight, Rob, Alison Vrbanac, Bryn C. Taylor, Alexander Aksenov, Chris Callewaert, Justine Debelius, Antonio Gonzalez, Tomasz Kosciolek, Liya-Iny McCall, Daniel McDonald, Alexey V. Melnik, Jamie T. Morton, Jose Navas, Thomas Quinn, Jon G. Sanders, Austin D. Swafford, Luke R. Thompson, Anupriya Tripathi, Zhenjiang Xu, Jesse R. Zaneveld, and Qiyun Zhu. 2018. ‘Best practices for analysing microbiomes’, Nature Reviews Microbiology, 16: 410–22.

Lanoil, Brian D., Roger Sassen, Myron T. La Duc, Scott T. Sweet, and Kenneth H. Nealson. 2001. ‘Bacteria and Archaea physically associated with Gulf of Mexico gas hydrates’, Applied and Environmental Microbiology, 67: 5143–53.

Lee, Michael D. 2019. ‘GToTree: a user-friendly workflow for phylogenomics’, Bioinformatics, 35: 4162–64.

Letunic, Ivica, and Peer Bork. 2021. ‘Interactive Tree Of Life (iTOL) v5: an online tool for phylogenetic tree display and annotation’, Nucleic Acids Research, 49: W293–W96.

Lever, Mark A., Kathryn L. Rogers, Karen G. Lloyd, Jörg Overmann, Bernhard Schink, Rudolf K. Thauer, Tori M. Hoehler, and Bo Barker Jørgensen. 2015. ‘Life under extreme energy limitation: a synthesis of laboratory- and field-based investigations’, FEMS Microbiology Reviews, 39: 688–728.

Lin, H., and Shyamal D. Peddada. 2020. ‘Analysis of compositions of microbiomes with bias correction’, Nature Communications, 11: 3514.

Lloyd, Karen G., L. Schreiber, Dorthe Groth Petersen, K. U. Kjeldsen, Mark A. Lever, Andrew D. Steen, Ramunas Stepanauskas, Michael Richter, Sara Kleindienst, Sabine Lenk, Andreas Schramm, and Bo Barker Jørgensen. 2013. ‘Predominant archaea in marine sediments degrade detrital proteins’, Nature, 496: 215–18.

McMurdie, P. J., and S. Holmes. 2013. ‘phyloseq: an R package for reproducible interactive analysis and graphics of microbiome census data’, PLoS One, 8: e61217.

Miller, Hannah E. 2014. ‘A 500-year sediment lake record of anthropogenic and climatic influence on water quality’, Environmental Science & Technology, 48: 4995–5004.

Mills, Katie. 2017. ‘Deciphering long-term records of natural variability and anthropogenic impacts in lake sediments’, Palaeogeography, Palaeoclimatology, Palaeoecology, 485: 1–16.

Morono, Y., T. Terada, J. Kallmeyer, and F. Inagaki. 2009. ‘Discriminative detection and enumeration of microbial life in marine subsurface sediments’, The ISME Journal, 3: 503□11.

Morono, Yuki, Akira Ijiri, Kenji Wakamatsu, and, et al. 2020. ‘Aerobic microbial life persists in oxic marine sediment as old as 101.5 million years’, Nature Communications, 11: 3626.

Nearing, Jacob T., Gavin M. Douglas, Molly G. Hayes, Jocelyn MacDonald, Dhwani K. Desai, Nicole Allward, Casey M. A. Jones, Robyn J. Wright, Akhilesh S. Dhanani, Andre M. Comeau, and Morgan G. I. Langille. 2022. ‘Microbiome differential abundance methods produce different results across 38 datasets’, Nature Communications, 13: 342.

Normark, William R., David J. W. Piper, Brian W. Romans, Jacob A. Covault, Peter Dartnell, and Ray W. Sliter. 2009. ‘Submarine canyon and fan systems of the California Continental Borderland.’ in, *Geological Society of America Special Paper* (Geological Society of America: Boulder, CO).

Nunoura, Takuro, Barbara Soffientino, Anna Blazejak, Jun Kakuta, Hiroshi Oida, Axel Schippers, and Ken Takai. 2009. ‘Subseafloor microbial communities associated with rapid turbidite deposition in the Gulf of Mexico continental slope (IODP Expedition 308)’, FEMS Microbiology Ecology, 69: 410–24.

Oksanen, Jari, Gavin L. Simpson, F. Guillaume Blanchet, Roeland Kindt, Pierre Legendre, Peter R. Minchin, R. B. O’Hara, Peter Solymos, M. Henry H. Stevens, Eduard Szoecs, and Helene Wagner. 2024. “vegan: Community Ecology Package.” In.: R Foundation for Statistical Computing.

Orcutt, Beth N., Jason B. Sylvan, Nina J. Knab, and Katrina J. Edwards. 2011. ‘Microbial ecology of the dark ocean above, at, and below the seafloor’, Microbiology and Molecular Biology Reviews, 75: 361–422.

Orsi, W.□D. 2018. ‘Ecology and evolution of seafloor and subseafloor microbial communities’, Nature Reviews Microbiology, 16: 671–83.

Orsi, W.□D., T. Magritsch, S. Vargas, Ö.□K. Coskun, A. Vuillemin, S. Höhna, G. Wörheide, S. D’Hondt, B.□J. Shapiro, and P. Carini. 2021. ‘Genome Evolution in Bacteria Isolated from Million□Year□Old Subseafloor Sediment’, mBio, 12: e01150□21.

Orsi, William D., Tobias Magritsch, Sergio Vargas, Ömer K. Coskun, Aurèle Vuillemin, Sebastian Höhna, Gert Wörheide, Steven D’Hondt, B. Jesse Shapiro, and Paul Carini. 2021. ‘Genome Evolution in Bacteria Isolated from Million-Year-Old Subseafloor Sediment’, mBio, 12.

Orsi, William D., Thomas A. Richards, and Warren R. Francis. 2018. ‘Predicted microbial secretomes and their target substrates in marine sediment’, Nature Microbiology, 3: 32–37.

Parada, A. E., D. M. Needham, and J. A. Fuhrman. 2016. ‘Every base matters: assessing small subunit rRNA primers for marine microbiomes with mock communities, time series and global field samples’, Environ Microbiol, 18: 1403–14.

Price, Morgan N., Paramvir S. Dehal, and Adam P. Arkin. 2010. ‘FastTree 2 – approximately maximum-likelihood trees for large alignments’, PLoS One, 5: e9490.

Prosser, James I., Brendan J. M. Bohannan, Thomas P. Curtis, Richard J. Ellis, Mary K. Firestone, Robert P. Freckleton, Jessica L. Green, Laurence E. Green, Ken Killham, Jay T. Lennon, A. Mark Osborn, Martin Solan, Christopher J. van der Gast, and J. Peter W. Young. 2007. ‘The role of ecological theory in microbial ecology’, Nature Reviews Microbiology, 5: 384–92.

Quast, C., E. Pruesse, P. Yilmaz, J. Gerken, T. Schweer, P. Yarza, J. Peplies, and F. O. Glockner. 2013. ‘The SILVA ribosomal RNA gene database project: improved data processing and web-based tools’, Nucleic Acids Res, 41: D590–6.

Ramírez, G.□A., D. Graham, and S. D’Hondt. 2018. ‘Influence of commercial DNA extraction kit choice on prokaryotic community metrics in marine sediment’, Limnology and Oceanography: Methods, 16: 525□36.

Ramírez, G. A., S. L. Jørgensen, R. Zhao, and S. D’Hondt. 2018. ‘Minimal Influence of Extracellular DNA on Molecular Surveys of Marine Sedimentary Communities’, Frontiers in Microbiology, 9: 2969.

Romans, B. W., W. R. Normark, M. M. McGann, and J. A. Covault. 2009. ‘Coarse-grained sediment delivery to deep-water basins’, Journal of Sedimentary Research, 79: 712–27.

Russell, Joseph A., Rosa León-Zayas, Kelly Wrighton, and Jennifer F. Biddle. 2016. ‘Deep Subsurface Life from North Pond: Enrichment, Isolation, Characterization and Genomes of Heterotrophic Bacteria’, Frontiers in Microbiology, 7: 678.

Shannon, Claude E., and Warren Weaver. 1949. The Mathematical Theory of Communication (University of Illinois Press: Urbana, IL).

Simon, Marie P. 2023. ‘Lake sediments as archives of short- and long-term environmental change’, Quaternary Research, 100: 1–14.

Spang, Anja, Jimmy H. W. Saw, Steffen L. Jørgensen, Katarzyna Zaremba-Niedzwiedzka, Joran Martijn, Anders E. Lind, Roel van Eijk, Christa Schleper, Lionel Guy, and Thijs J. G. Ettema. 2015. ‘Complex archaea that bridge the gap between prokaryotes and eukaryotes’, Nature, 521: 173–79.

Sperfeld, Michaela, Katharina Zecher, Daniela Liebschner, Thomas Riedel, Simon Schulz, Meinhard Simon, and Thorsten Brinkhoff. 2024. ‘Algal methylated compounds shorten the lag phase of Phaeobacter inhibens bacteria’, Nature Microbiology, 9: 190–201.

Sprouffske, Kathleen, and Andreas Wagner. 2016. ‘Growthcurver: an R package for obtaining interpretable metrics from microbial growth curves’, BMC Bioinformatics, 17: 172.

Stamatakis, Alexandros. 2014. ‘RAxML version 8: a tool for phylogenetic analysis and post-analysis of large phylogenies’, Bioinformatics, 30: 1312–13.

Valette-Silver, Nathalie J. 1993. ‘The use of sediment cores to reconstruct historical trends in contamination of estuarine and coastal sediments’, Environmental Science & Technology, 27: 13–20.

Vigneron, Adrien, Perrine Cruaud, Patricia Pignet, Jean-Claude Caprais, Marie-Anne Cambon-Bonavita, Anne Godfroy, and Laurent Toffin. 2013. ‘Archaeal and anaerobic methane oxidizer communities in the Sonora Margin cold seeps, Guaymas Basin (Gulf of California)’, The ISME Journal, 7: 1595–608.

Vuillemin, A., S. Vargas, Ö.□K. Coskun, R. Pockalny, R.□W. Murray, D.□C. Smith, S. D’Hondt, and W.□D. Orsi. 2020. ‘Atribacteria Reproducing over Millions of Years in the Atlantic Abyssal Subseafloor’, mBio, 11: e01937–20.

Waite, David W., Maria Chuvochina, Christoph Pelikan, Donovan H. Parks, Pelin Yilmaz, Michael Wagner, Alexander Loy, Takeshi Naganuma, Ryuhei Nakai, William B. Whitman, and Philip Hugenholtz. 2020. ‘Proposal to reclassify the proteobacterial classes Deltaproteobacteria, Oligoflexia, and Thermodesulfobacteria into four phyla reflecting major functional capabilities’, International Journal of Systematic and Evolutionary Microbiology, 70: 5972.

Walsh, Emily A., John B. Kirkpatrick, Scott D. Rutherford, David C. Smith, Mitchell Sogin, and Steven D’Hondt. 2016. ‘Bacterial diversity and community composition from seasurface to subseafloor’, The ISME Journal, 10: 979–89.

Warrick, J. A., and K. L. Farnsworth. 2009. ‘Sources of sediment to the coastal waters of the Southern California Bight’, Geological Society of America Special Papers, 454: 39–52.

Xiao, Guangnian, Tian Wang, Xinqiang Chen, and Lizhen Zhou. 2022. ‘Evaluation of Ship Pollutant Emissions in the Ports of Los Angeles and Long Beach’, Journal of Marine Science and Engineering, 10: 1206.

Yang, Yi, Yaozhi Zhang, Natalie L. Cápiro, and Jun Yan. 2020. ‘Genomic characteristics distinguish geographically distributed Dehalococcoidia’, Frontiers in Microbiology, 11: 546063.

Yin, X., A. Zhu, Y. Li, and S. Wang. 2021. ‘Subgroup level differences of physiological activities in Lokiarchaeota’, The ISME Journal, 15: 2051–64.

